# *In vivo* Quantification of White Matter Pathways in the Human Hippocampus

**DOI:** 10.1101/2025.01.13.632268

**Authors:** Melisa Gumus, Athanasios Bourganos, Michael L. Mack

**Author notes:** Corresponding author:Melisa Gumus, Department of Psychology, University of Toronto 100 St. George Street, Toronto, ON M5S 3G3, Canada.

## Abstract

The hippocampus is a key structure in cognition. Although much research has focused on defining the functions of its anatomically distinct subfields, the communication among these subfields within the hippocampal circuit, supported by white matter pathways, is theoretically key to emergent cognitive function. Yet, hippocampal white matter connections in humans have not been fully explored *in vivo*. By leveraging diffusion weighted imaging and a large healthy sample (N=831), we developed a processing pipeline for *in vivo* quantification of human hippocampal pathways. We provided evidence for monosynaptic and trisynaptic pathway-related connections in humans, supporting the described hippocampal circuit in *ex vivo* and animal studies. In addition to hemispheric and sex differences, the individual variability in hippocampal pathways was linked to cognitive abilities. Thus, *in vivo* characterization of human hippocampal pathways highlights the individual differences within these structures and paves the way for their implications in cognition, aging, and diseases.

**Conflict of interest:** The authors declare no conflicts of interest.

## 1. Introduction

The hippocampus is a critical brain region in supporting wide range of cognitive functions including episodic memory, category learning, spatial navigation and decision making (Eichenbaum, 2004; Mack et al., 2018; Schapiro et al., 2017; Scoville & Milner, 1957; Shohamy & Turk-Browne, 2013). With its extensive involvement in cognition, it is also implicated in variety of medical conditions from neurodegenerative to psychiatric disorders (de Flores et al., 2015; Foo et al., 2017; Maruszak & Thuret, 2014). Histological and cytoarchitectural evidence highlights anatomically distinct hippocampal subfields such as dentate gyrus (DG), cornu ammonis area 1 (CA1), CA2/3/4, and subiculum (SUB) as well as entorhinal cortex (ERC) as the major input/output region (Duvernoy et al., 2013). These structures are recruited during many cognitive functions (Aimone et al., 2011; Leutgeb et al., 2007; O’Reilly et al., 2014) as information travels within the hippocampal pathways connecting the subfields and ERC (Anand & Dhikav, 2012). Yet, there is limited empirical *in vivo* characterization of human hippocampal white matter connections, which is fundamental to understanding the role of hippocampus in cognition and its implications in disorders (Karat et al., 2023; Yassa et al., 2010, 2011; Zeineh et al., 2001, 2017).

Our knowledge regarding the hippocampal circuit is mostly inferred from animal studies (David & Pierre, 2006; Patten et al., 2015), which motivates the theoretical framework for the flow of information within the hippocampal circuit (Fig. 1a) (O’Reilly et al., 2014; Schapiro et al., 2017). There are two main pathways linking the hippocampal subfields and ERC: trisynaptic (TSP) and monosynaptic (MSP) (Patten et al., 2015). ERC is the main input of the hippocampal circuit, and thus of both pathways (David & Pierre, 2006; Witter et al., 2017). As part of TSP, ERC projects its unidirectional connections onto DG through the perforant pathway (Patten et al., 2015; Witter et al., 2017). Unmyelinated mossy fibers then link DG to CA3, which then connects to CA1 via schaffer collaterals. MSP, on the other hand, is composed only of ERC connections that directly project onto CA1 without stopping at DG (Yeckel & Berger, 1995). To complete the hippocampal circuit, CA1 connects to SUB, which is then linked back to ERC (Patten et al., 2015). Besides ERC, SUB is also considered as an output from the hippocampal circuit, delivering processed information to cortical and subcortical regions although its specific functions in cognition remain elusive (Roy et al., 2017). Post-mortem work points to a high resemblance of the described hippocampal circuit in humans (Beaujoin et al., 2018). However, the challenge has been to investigate these structures in humans with an *in vivo* design.

**Fig. 1:**
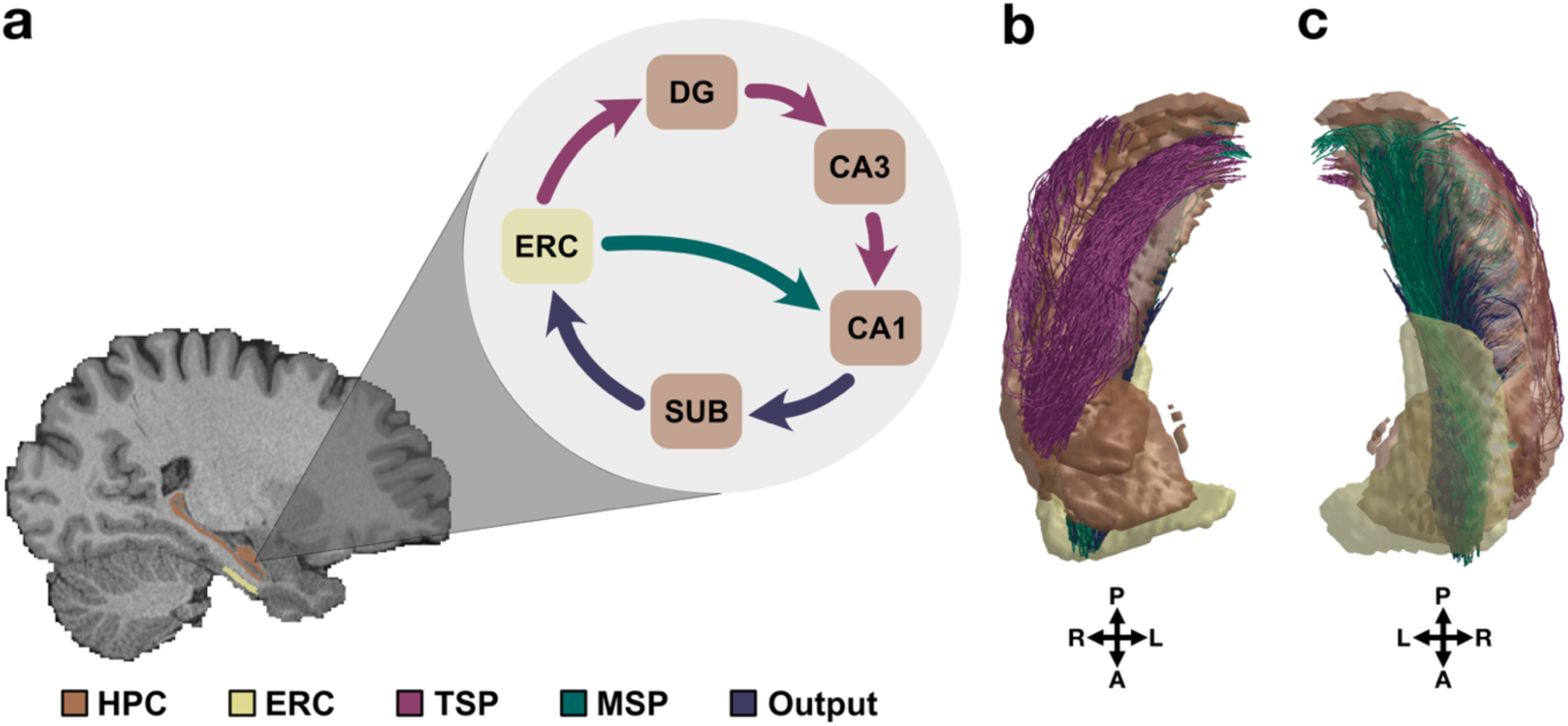
Theorized hippocampal circuit and *in vivo* quantification of its connections in humans. a, Hippocampal circuit links the subfields of the hippocampus and ERC. MSP connects ERC to CA1 while TSP spans across ERC, DG, CA3 and CA1. We refer to the CA1-SUB and SUB-ERC as output connections since they complete the hippocampal circuit. **b,** Quantified TSP-related connections in humans are visible from superior view. **c)** MSP-related and output connections in humans are observed from the inferior view.

Developments in Magnetic Resonance Imaging (MRI) techniques within the last two decades enabled *in vivo* measurements of the human hippocampal subfields (DeKraker et al., 2021; Iglesias et al., 2015; Mueller et al., 2007; Van Leemput et al., 2009; Yushkevich et al., 2009). Extensive description of the anatomical distinctions between the hippocampal subfields advanced research on their involvement in cognition. (Córdova et al., 2019; Duncan et al., 2012; Suthana et al., 2015). For example, human CA1 integrates new experiences with the existing memories that have overlapping features (Schlichting et al., 2014) while CA3 and DG separate certain experiences and store them as distinct representations (Bakker et al., 2008; Lacy et al., 2011). One missing piece of the puzzle in the hippocampal circuit is understanding the role of hippocampal pathways connecting these subregions. Although the neuroanatomy of the hippocampal formation is largely consistent across species such as monkeys, rats and humans, there exist substantial structural differences in the organization of its connections with itself and cortical areas (Clark & Squire, 2013; David & Pierre, 2006; Zeineh et al., 2017). Thus, *in vivo* approaches are key to characterize the cognitive relevance of white matter pathways of the human hippocampal circuit.

Diffusion imaging is the dominant technique for *in vivo* evaluation of white matter connections in the human brain, revealing insights into their structure and the underlying cognitive processes (Le Bihan & Johansen-Berg, 2012). The particular imaging relies on Brownian motion of water molecules which diffuse more freely in the direction of fibers, allowing differentiation of white matter structures (Le Bihan & Johansen-Berg, 2012). Although the hippocampus is a relatively small region, novel diffusion processing methods have enabled researchers to investigate broader hippocampal connections with cortical and subcortical regions in humans (Dalton et al., 2022; Huang et al., 2021; Maller et al., 2019). Distinct cortical regions, including medial temporal lobe, parietal and occipital regions, are preferentially connected along the anterior-posterior axis of the human hippocampus (Dalton et al., 2022). *In vivo* neuroimaging methods also infer measures of neuroplasticity within the human hippocampus such as remodelling of white matter connections during learning, even within short time intervals (Sagi et al., 2012). Such tools are also feasible in investigating aging related changes within the human hippocampal circuit. For example, white matter connections between the hippocampal subfields present reduced structural integrity in older adults (Elsaid et al., 2022) and the degradation of perforant connections relates to memory deficits (Yassa et al., 2011). Novel *in vivo* neuroimaging techniques and processing pipelines specific to the human hippocampus is critical in understanding the role of hippocampal circuit in cognition. Thus, it is necessary to characterize *in vivo* hippocampal white matter connections in a large sample to establish a normative model of human hippocampal circuit and the individual variability among its interconnections.

Although diffusion techniques are powerful in *in vivo* investigations, certain limitations need to be addressed (Schilling et al., 2019). The lack of sensitive processing methods and the difficulty with their validation and reproducibility could pose a challenge to characterizing the human hippocampal pathways. To increase transparency and reproducibility, we provide a detailed protocol and documentation on the processing pipeline for hippocampal white matter connections. The large sample size included in this study improves the generalizability of the results. There have also been limitations around image resolution and the complex microstructure of white matter pathways such as crossing fibers. However, innovative approaches in diffusion MRI allow for more sensitive and detailed examination of white matter connections in humans (Fan et al., 2016; McNab et al., 2013) with the ability to evaluate fiber orientations within a voxel and reconstruct tractography with high resolution (Basser et al., 2000; Wedeen et al., 2012). These developments allow for white matter connections to be estimated within the brain regions with complex structural organization, such as the hippocampus. Indeed, such algorithms previously revealed the hippocampal connections with the rest of the brain (Dalton et al., 2022; Huang et al., 2021). However, there has not been an extensive *in vivo* investigation, specifically into the white matter connections among the hippocampal subfields and ERC in humans, considering the theorized pathways (i.e., TSP and MSP).

We extended the existing *in vivo* diffusion efforts (Dalton et al., 2022; Elsaid et al., 2022; Huang et al., 2021; Karat et al., 2023) to the white matter connections of the hippocampal circuit in humans by leveraging a large sample of healthy young adults. By utilizing the recent advancements in diffusion imaging, we quantified hippocampal white matter pathways in humans, including MSP-, TSP-related and output connections (Fig. 1b, 1c). This work supported the theorized anatomical structures of the hippocampal circuit derived from *ex vivo* and animal model studies. The individual variability we observed in the hippocampal pathways was also related to cognition. Thus, we present *in vivo* evidence for the feasibility of mapping the white matter connections of the hippocampal circuit in humans, using diffusion imaging. To enable further investigation of hippocampal pathways and their role in cognition, aging and diseases, the diffusion processing pipeling and all reported data are available as a public resource with detailed documentation.

## 2. Results

2.1. *In vivo* Reconstruction of Hippocampal White Matter Connections in Humans

The developed pipeline demonstrated the feasibility of *in vivo* quantification of hippocampal white matter connections in humans with diffusion weighted imaging. We first generated whole brain tractographies at the participant level. These tractographies were then masked by the segmentation images, that included the hippocampal subfields and ERC, to isolate the white matter connections specific to the hippocampal circuit. Similar processing was also performed on the template diffusion image that was generated based on 831 participants (Fig. 2a).

**Fig. 2:**
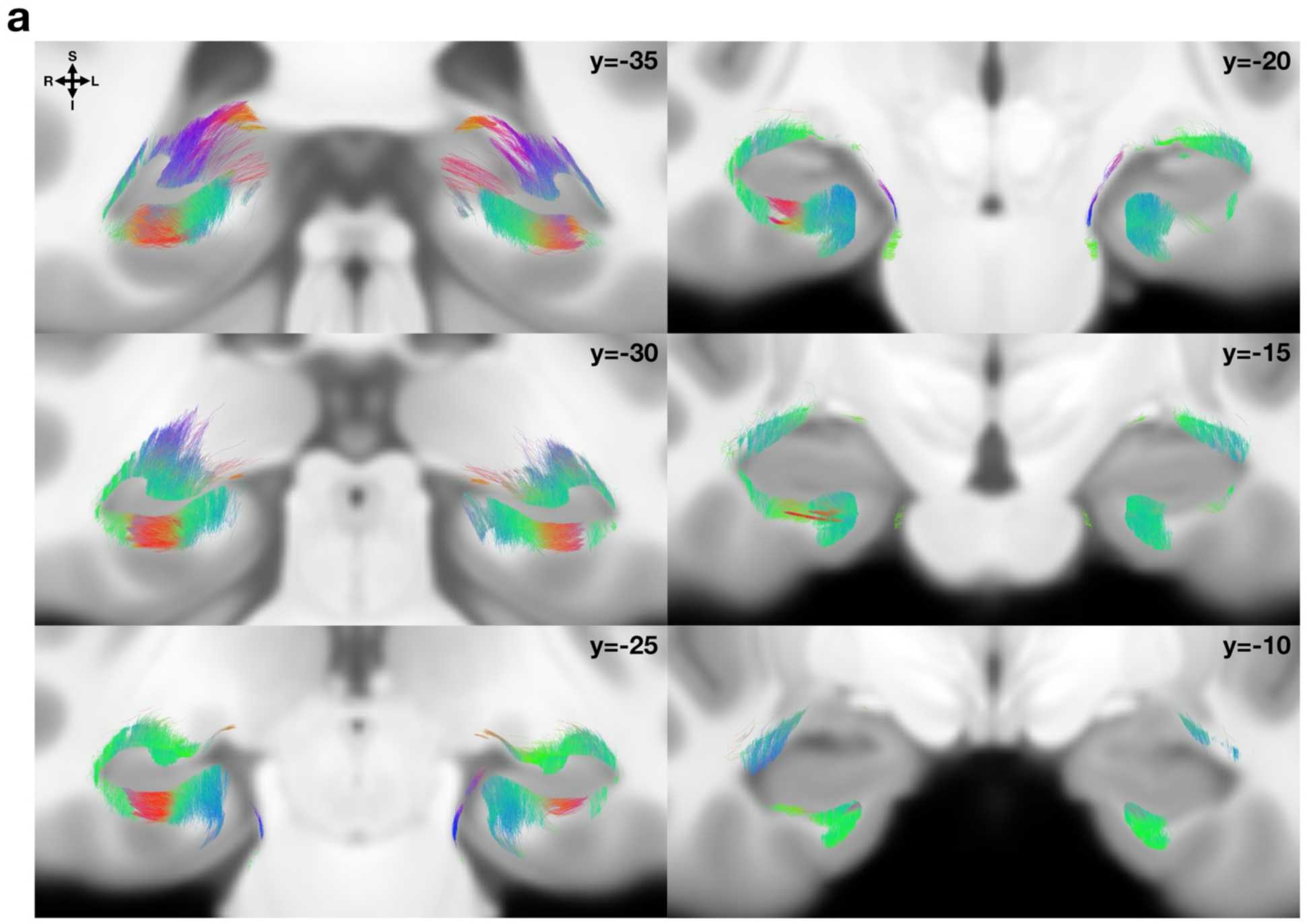
***In vivo* reconstruction of white matter connections in the human hippocampus. a,** White matter connections in the human hippocampus extend from anterior to posterior end. The visualized connections include those between hippocampal subfields, ERC as well as the self-connections. The depicted connections are based on the template image that includes 831 participants. White matter connections follow the standard color-coding of tractography: red for right-left, green for anterior-posterior, and blue for dorsal-ventral. For oblique orientations, the colours would be blended; combination of red and blue orientations yields magenta.

Diffusion imaging cannot reveal the directionality of the connections such as those from ERC to CA1; however, it can provide insight into the overall alignment of the connections. The reconstructed hippocampal white matter connections in humans extended along the anterior-posterior axis of the hippocampus, linking the subfields and ERC (Fig. 2a), consistent with animal and *ex vivo* studies (Andersen, 2006; Zeineh et al., 2017). Based purely on the quantified connections, hippocampal structure was readily distinguished in coronal slices. Although most connections were in the anterior-posterior direction (i.e., depicted in green), certain inferior connections lied along the right-left axis (i.e., depicted in red) of the hippocampus. Additionally, towards the posterior hippocampus, some connections were positioned along the inferior-superior extent, possibly forming into larger efferent white matter connections of the hippocampus.

### 2.2. White Matter Connections Isolated for Specific Hippocampal Pathways in Humans

The quantified connections in the human hippocampus were then isolated for the traditionally defined hippocampal pathways, namely MSP-, TSP-related and output connections. MSP-related connections were specific to those between ERC and CA1. These connections converged onto a relatively specific region of ERC but exhibited a wider connectivity on CA1, curving laterally towards the ends (Fig. 3a). The sparse CA1 connectivity might be arising from multiple factors such as the non-uniform spatial input profile CA1 (Druckmann et al., 2014) or scattered patterns of certain CA1 cell types (Bocchio et al., 2020).

**Fig. 3:**
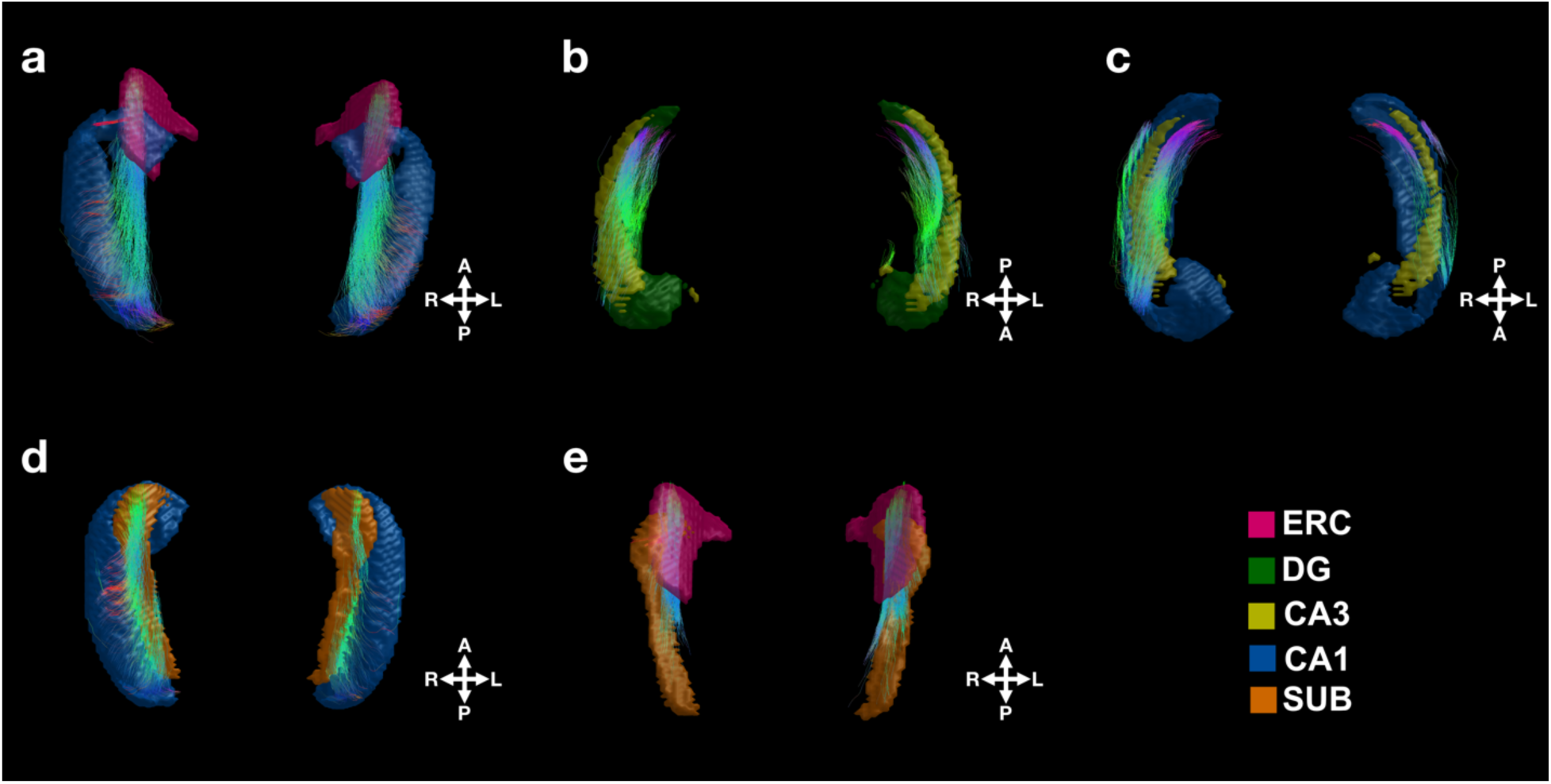
White matter connections of theorized hippocampal pathways in humans. The quantified human hippocampal white matter connections were isolated for traditionally defined hippocampal pathways. The population template of hippocampal connections is depicted. **a,** MSP-related connections link ERC and CA1. **b,** DG-CA3 and **c,** CA3-CA1 connections are part of the TSP-related connections. **d,** CA1-SUB and **e,** SUB-ERC are referred to as the output connections of the hippocampal circuit.

On the other hand, TSP-related connections presented a relatively slimer pattern (Fig. 3b, 3c). These included 2 sets of connections: DG-CA3 (Fig. 3b) and CA3-CA1 (Fig. 3c). Both TSP-related connections (i.e., DG-CA3 and CA3-CA1) appeared as bundles of relatively shorter-range connections attaching corresponding 2 subfields along the long-axis of the hippocampus. TSP-related connections were positioned more superior to the hippocampus while those of MSP-related were more inferior. For the rest of the analyses, TSP-related connections were masked by relatively larger white matter structures of the hippocampus, specifically fornix, fimbria and mammillary bodies to avoid any potential overlap as they also lie superior to the hippocampus. Traditionally, TSP also includes connections between ERC and DG. However, these connections did not yield high quality reconstruction in the current work, thus, were excluded from the rest of the analyses. The difficulty in visualizing ERC-DG connections is due the specific curvature of DG as reported in previous *ex vivo* diffusion work (Augustinack et al., 2010).

Lastly, we isolated 2 output connections: CA1-SUB (Fig. 3d) and SUB-ERC (Fig. 3e). We refer to these as output connections since they complete the hippocampal circuit. CA1-SUB connections resembled MSP-related connections, landing on a widespread region of CA1 with curved ends. Both ERC-CA1 and CA1-SUB presented long-distance connections, consistent with their description in post-mortem work (Modo et al., 2023).

### 2.3. White Matter Connections are Differentially Distributed in the Hippocampal Circuit

Quantifying participant-specific hippocampal white matter connections uncovered differential distribution of human hippocampal pathways. Based on 10 million streamlines in each participant, which was down sampled to 2 million for biologically plausible connections, the hippocampal circuit had a mean of 2665 ± 516 streamlines with individual differences ranging from 1210 to 4561 streamlines (Fig. 4a). Averaging the streamline densities across 831 participants, we observed that the connections were differentially distributed across the traditionally defined hippocampal pathways (Fig. 4b). Only about 5% of all hippocampal streamlines were within the ERC-CA1 connections, making MSP-related connections the smallest in the hippocampal circuit. Both TSP-related connections included twice as many streamlines as MSP-related connections. Specifically, 14% of streamlines was part of DG-CA3 and about 13% was within CA3-CA1 connections, bringing TSP-related connections to 27%. Most of the streamlines within the hippocampal circuit fell within the output connections: 26% of all hippocampal streamlines was part of CA1-SUB and 23% was within the SUB-ERC connections.

**Fig. 4:**
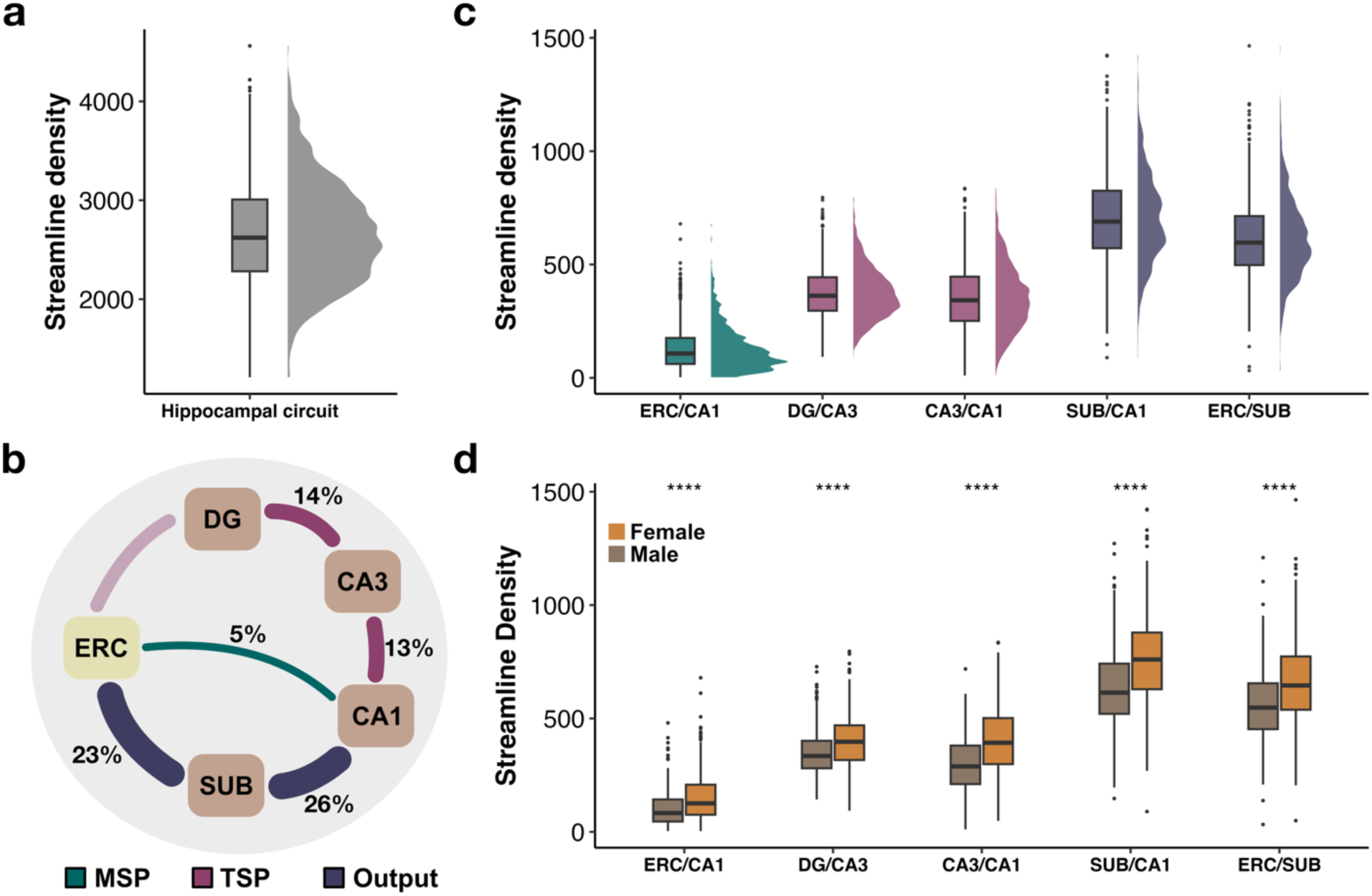
Individual variability in the differentially distributed hippocampal white matter connections. a, Streamline density distribution in human hippocampal circuit. **b,** Hippocampal white matter connections are differentially distributed across hippocampal pathways. Percentages represent the distribution of streamlines that belong to a particular connection between 2 regions. **c,** Streamline density of hippocampal white matter connections, plotted across the traditional defined hippocampal pathways. Output connections present the greatest streamline density, which is followed by TSP-related and then MSP-related connections. **d,** Streamline density of the hippocampal pathways present biological sex differences with greater streamline density in females than males.

### 2.3. Individual Variability in the Human Hippocampal Pathways

In addition to assessing the distribution of streamlines across the hippocampal pathways, we evaluated participant-specific hippocampal pathways. Analyses of Variance revealed statistically significant differences between the streamline density of the 5 hippocampal pathways, F (5, 4148) = 1951, *p*=2E-16 (Fig. 4c). The streamline density of TSP-related connections, both DG- CA3 and CA3-CA1, was statistically greater than the MSP connections (i.e., ERC-CA1), p=3.34E-08, 95% CI = [225.42, 265.27] and p=3.34E-08, 95% CI = [203.34, 243.19] (Fig. 4c).

There was no significant difference between the TSP-related connections (i.e., DG-CA3 and CA3-CA1). Streamline density of both output connections, CA1-SUB and SUB-ERC, was significantly greater than CA3-CA1 of TSP-related connections, p=3.34E-08, 95% CI = [330.96, 370.81] and p=3.34E-08, 95% CI = [241.21, 281.07] (Fig. 4c). Similarly, output connections, CA1-SUB and SUB-ERC, showed significantly larger streamline density than DG-CA3 connections, p=3.34E-08, 95% CI = [308.88, 348.73] and p=3.34E-08, 95% CI = [219.14, 258.99].

The individual variability in streamline densities also differed across the 5 hippocampal pathways (Fig. 4c, inset violin plots). MSP-related connections exhibited a very high variability in streamline density with a coefficient of variation of 74.20%, indicating a wide dispersion around the mean. Some of this variability might be driven by its skewed distribution (Fig. 4c, inset violin plots). Although these connections are not zero, they are relatively small. These streamline density estimates are based 10 million streamlines generated within the whole brain, which is a standard practice. Studies with small or medium sample sizes may benefit from generation of more than 10 million streamlines. However, this was not computationally feasible in the current study with such a large sample size. CA3-CA1 of TSP-related connections presented the second greatest individual variability following MSP-related connections with a coefficient of variation of 40.02%. Lastly, DG-CA3 of TSP-related connections as well as the output connections, SUB-CA1 and CA1-ERC, included moderate variability of streamline density with a coefficient of variation of 29.78%, 27.62% and 28.75%.

### 2.4. Hemispheric Differences in the Human Hippocampal Pathways

We observed hemispheric differences in streamline density of MSP-, TSP-related, and output connections (Extended Data Fig. 1a). Pairwise comparisons revealed greater streamline density of MSP-related connections (i.e., ERC-CA1) in the left hippocampus than the right, t(830) = 4.87, p = 1.36E-06. On the contrary, the rest of the hippocampal pathways presented larger streamline density in the right hippocampus than the left. Specifically, right TSP-related connections, DG-CA3 and CA3-CA1, had greater streamline density than the left, t(829) = 4.41, p = 1.16E-05 and t(829) = 40.52, p<2.2E-16. Similarly, output connections, CA1-SUB and SUB- ERC, in the right hippocampus included significantly larger streamline density than the left hippocampus, t(830) = 38.48, p < 2.2E-16 and t(830) = 12.90, p<2.2E-16. All hemispheric differences remained significant after the Bonferroni correction for multiple comparisons at the alpha level of 0.01 (0.05/5 comparisons). In addition, we investigated whether the individual variability or the hemispheric difference of the hippocampal pathways were driven by inequal volumes of ERC and the hippocampal subfields. We corrected the streamline density of hippocampal connections by the volume of the 2 regions each connection links to. Above comparisons remained significant, indicating no volume effects on our results (Extended Data Fig. 1b).

### 2.5. Sex Differences in the Human Hippocampal Pathways

All quantified human hippocampal pathways (i.e., MSP-, TSP-related, and output connections) demonstrated biological sex differences with greater streamline density in females than males (Fig. 4d). Specifically, females presented significantly greater streamline density in MSP-related connections (i.e., ERC-CA1) t(810.08) = 7.65, p = 5.9E-14. Females also had greater streamline density in both TSP-related connections (i.e., DG-CA3 and CA3-CA1), t(827.91) = 7.42, p =2.9E-13 and t(825.47) = 11.94, p < 2.2E-16. Lastly, both output connections, CA1-SUB and SUB-ERC, also had significantly more streamline density in females than males, t(824.45) = 10.04, p < 2.2E-16 and t(828.46) = 8.49, p < 2.2E-16.

### 2.6. Individual Variability in Trisynaptic Connections Relates to Cognition

Although proportion of TSP-related connections (i.e., 27%) is relatively higher than that of MSP (i.e., 5%), TSP-related connections are pooled from 2 separate connections, namely DG-CA3 and CA3-CA1. Given the importance of TSP in cognitive processing, we further explored the connectivity structure within the TSP subnetwork by estimating the connection probabilities from the perspective of each of its subfields. For example, if we consider all hippocampal connections that land on CA3, 42% of them were linked to DG and 39% of CA3 connections landed on CA1, highlighting dense connectivity within the TSP subnetwork (Fig. 5a). Similarly, 51% of CA1 connections linked to SUB to form the output connections, and about 26% of CA1 connections were shared with CA3 (Fig. 5b). The relative proportion of hippocampal connections within the TSP subnetwork maps out a clear pathway consistent with the theorized structure of TSP (Andersen et al., 1971; David & Pierre, 2006).

**Fig. 5:**
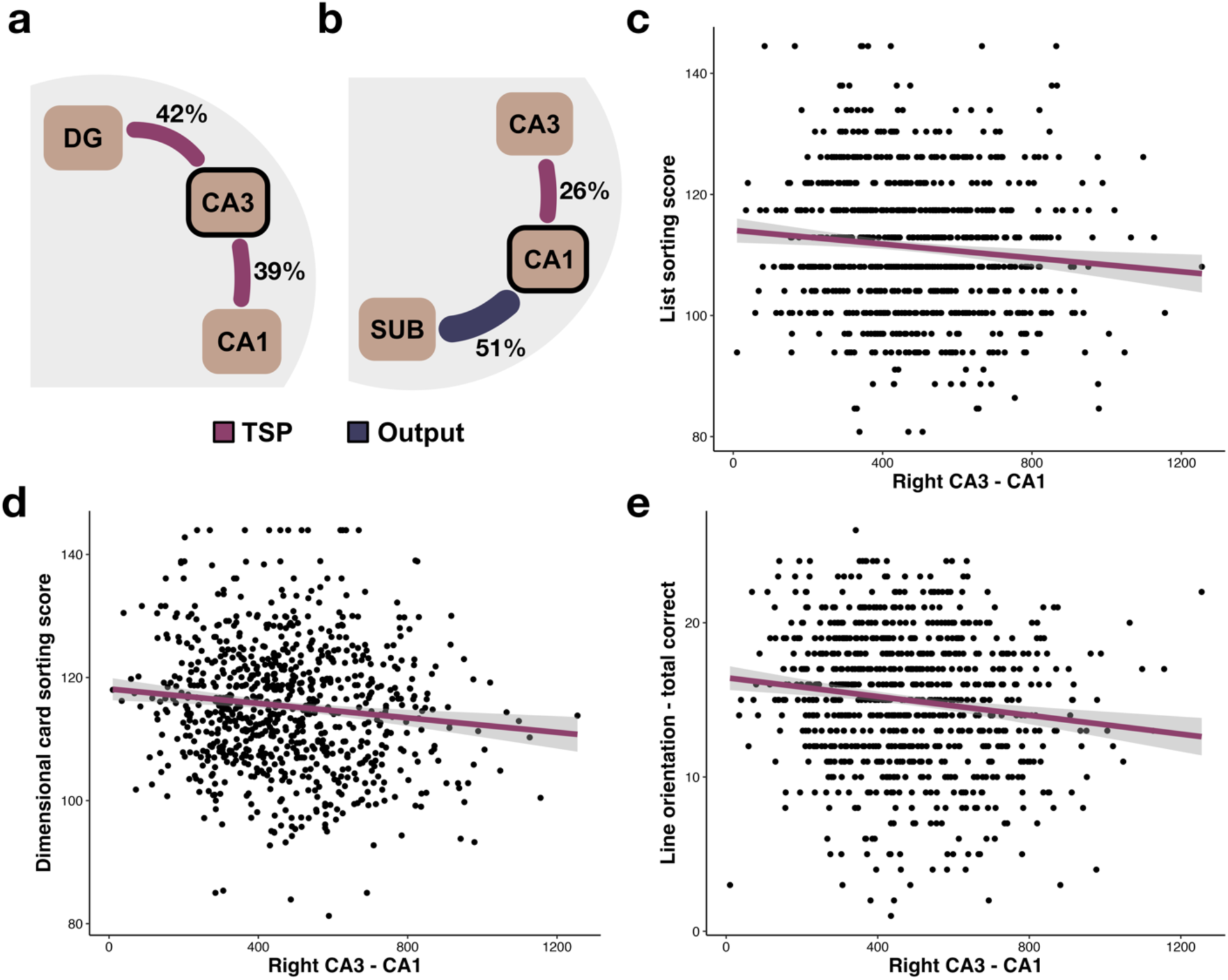
Individual variability in trisynaptic pathway-related connections is linked to cognition. a, CA3 is densely connected to DG and CA1 within the TSP-subnetwork. **b,** Most of CA1 connections are shared with SUB, but significant percentage of its connections are linked to CA3. Individual variability in the streamline density of right CA3-CA1 of TSP-related connections is associated with **c,** list sorting scores (i.e., working memory), **d,** dimensional card sorting ability (i.e., cognitive flexibility), and **e,** line orientation performance (i.e., spatial orientation).

We also investigated whether streamline densities of the quantified hippocampal pathways (MSP-, TSP-related and output connections) were linked to cognition. Linear mixed effects models with age and sex covariance were fit to hippocampal pathways to predict cognitive scores. We observed that number of cognitive domains including working memory, spatial orientation and cognitive flexibility were related to the hippocampal pathways, specifically to TSP-related connections (i.e., CA3-CA1) (Fig. 5c-e). The streamline density of right CA3-CA1 connections were negatively associated with list sorting scores (i.e., working memory; β=-1.34, 95% CI [-2.35, -0.34], p = 0.009; Fig. 5c). Although unintuitive why reduced streamline density might be related to better working memory capacity, this is consistent with the importance of sparse TSP-like representations in learning and memory processes (Knierim & Neunuebel, 2016; Leutgeb et al., 2007; Niibori et al., 2012). Similarly, reduced streamline density of right CA3- CA1 was also associated with better performance in dimensional card sorting (i.e., cognitive flexibility; β=-1.75, 95% CI [-2.86, -0.63], p = 0.002; Fig. 5d) and line orientation total correct (i.e., spatial orientation; β=-3.75, 95% CI [-6.35, -1.16], p = 0.005; Fig. 5e). These results remained significant after multiple comparisons corrections with the Bonferroni test. We also observed that the streamline density of right SUB-CA1 was associated with verbal episodic memory scores (β=5.11, 95% CI [1.09, 9.13], p =0.013); however, this did not survive the Bonferroni correction.

## 3. Discussion

We provide *in vivo* evidence for the hippocampal pathways in humans, which support the neurobiological description of the hippocampal circuit developed by animal models and theoretical work (Andersen, 2006; Zeineh et al., 2017). We demonstrated the interconnectedness of the hippocampal subfields and ERC in humans as white matter connections extended along the long axis of the hippocampus. The traditionally defined hippocampal pathways, including MSP-, TSP-related and output connections, emerged from the quantified white matter connections, verifying the feasibility of diffusion methods for *in vivo* investigation of human hippocampal circuit. We observed that human hippocampal white matter connections were differentially distributed across the hippocampal pathways; TSP-related connections comprised more streamlines than the MSP-related connections but less than the output connections.

Moreover, characterizing hippocampal white matter connections at the participant level revealed the individual differences. Biological sex and hemisphere were only two sources of the variability. Such diversity observed in the human hippocampal pathways was fundamentally linked to cognitive abilities. Thus, *in vivo* quantification of human hippocampal pathways is critical in understanding the role of hippocampal circuit in cognition but its potential implications in aging and diseases.

Quantified white matter connections extended along the long axis of the human hippocampus, confirming the anatomy reported in *ex* vivo and animal studies (Andersen, 2006; Zeineh et al., 2017). Sorting these white matter connections according to each pair of hippocampal subfields and ERC uncovered the traditionally defined hippocampal pathways in humans, including MSP-, TSP-related and output connections. TSP-related connections included twice as more streamlines than those of MSP-related. This was consistent with *ex vivo* work; TSP consisted of dense connections while MSP connections were sparser (Johnston & Amaral, 2004), which was also implemented in the computational models of the hippocampus (Ketz et al., 2013; Schapiro et al., 2017). Despite fundamental differences between MSP and TSP, CA1 receives input from both ERC and CA3. We observed no overlap between these connections such that MSP-related connections landed mostly on inferior CA1 while TSP-related connections were more superior. Theoretical work suggests that both MSP and TSP feed into the output connections, including CA1-SUB and SUB-ERC, which complete the hippocampal circuit.

Large percentage of quantified hippocampal connections were part of the output connections, consistent with rodent work that CA1, SUB and ERC are heavily interconnected (Naber et al., 2001; Tamamaki & Nojyo, 1995). In addition to connections from CA1 to SUB, tracing studies in rodents revealed backward projections from SUB to CA1 (Xu et al., 2016). Future research should investigate whether the strong connections we observed between CA1-SUB arise from this bidirectional connectivity. On the other hand, SUB-ERC connections are considered a major output of the hippocampal circuit, allowing communication with subcortical and cortical region (David & Pierre, 2006; Witter et al., 2017). Dense SUB-ERC connections might be necessary for this important function. Previously, strong SUB-ERC connections were also reported across different species including rodents (Cembrowski et al., 2018), cats (Ino et al., 2001), and rabbits (Honda & Shibata, 2017) as well as in *ex vivo* human hippocampus (Beaujoin et al., 2018). Thus, we extended *ex vivo* findings to *in vivo* work using diffusion imaging, confirming the organization and characteristics of hippocampal circuit in humans. (Augustinack et al., 2010; Beaujoin et al., 2018; Ly et al., 2020).

*In vivo* quantification of human hippocampal pathways revealed great variability across individuals. Structural variance within the hippocampus is often related to aging and cognitive performance (Patel et al., 2020). We found that the hippocampal circuit presented hemispheric differences; TSP-related and output connections included more streamlines in the right hippocampus, while MSP-related connections were more prominent in the left hippocampus. These characteristics of the human hippocampal pathways could have implications in cognition, aging and diseases. For instance, recent diffusion work reported reduced structural integrity in the hippocampal connections in older adults, with more pronounced aging effects in the left

hippocampus (Elsaid et al., 2022). Considering our results that left hippocampus includes fewer streamlines in most pathways, the question becomes whether aging differentially influences the right versus left hippocampal circuit. Such structural variability in the hippocampal circuit likely arises during development, shaping cognitive abilities throughout the lifespan, as white matter pathways mature at different rates from early childhood to adulthood (Lebel et al., 2019). As one specific example, the differences in episodic memory performance in early childhood can be predicted by the white matter connections between the hippocampus and parietal lobe (Ngo et al., 2017). Our processing pipeline can further our understanding of the developmental trajectories of the hippocampal pathways and their implications throughout the lifespan and in aging.

Describing individual differences of the hippocampal white matter pathways is also impactful in predicting disease outcomes. One factor that contributes to the individual differences in the human brain is biological sex. Our results revealed that females exhibited significantly greater streamline density than males. Sex is, in fact, a significant contributor to the cognitive abilities, brain structure and the relationship between brain and cognition (Gur et al., 1999; Haier et al., 2005; Halpern, 2013). Clinical work should balance sex distributions in order to avoid potential confounds. Distinct subfields of the hippocampal circuit are suggested to be differentially vulnerable to various disorders (Small et al., 2011). For example, major depression patients were reported with reduced structural connectivity within DG (Rutland et al., 2019). As a result, white matter connections between the hippocampal subfields could also be distinctly implicated in diseases. Individual differences in the hippocampal circuit can relate to severity of disease symptomology. Understanding the variability in the hippocampal circuit, including the biological sex differences, can optimize the search for personalized treatments in diseases.

The individual differences in the human hippocampal pathways contributed to the diversity in cognitive abilities. We found that the variability in TSP-related connections was associated with several cognitive functions including working memory, spatial orientation ability and cognitive flexibility. With its critical role in declarative memory formation, some consider TSP as the backbone of information processing in the hippocampus (Zorumski, 2011). We previously reported the importance of TSP in learning exceptional items that need to be distinguished from similar items in memory (Schlichting et al., 2021). This ability to discriminate similar experiences was reduced in older adults, which was linked to age related structural degradation of TSP like connections (Yassa et al., 2010). Although the cognitive assessments available in this current study did not relate to MSP-like connections, in theory, various cognitive functions are differentially supported by the hippocampal pathways. *In vivo* quantification of human hippocampal pathways creates an opportunity to further investigate the contributions of MSP and TSP in cognition. Indeed, computational work posits distinct functions of hippocampal pathways; TSP rapidly binds elements of a specific experience into distinct memory traces, and MSP slowly builds unitized representations over many experiences (Schapiro et al., 2017). We introduce an analytical processing resource that can provide empirical evidence for the prominent theories.

Recent developments in diffusion techniques accelerated the *in vivo* characterization of hippocampal connections with subcortical and cortical regions (Dalton et al., 2022; Elsaid et al., 2022; Maller et al., 2019; Treit et al., 2018). For instance, systematic evaluation of diffusion imaging revealed that the hippocampus was much more connected to cortical regions than it was previously thought, providing the first *in vivo* evidence for hippocampal-cortical white matter connections in humans (Maller et al., 2019). Higher angular and diffusing sampling resolution enabled the detection of hippocampal connections with cingulate, that was not possible in previous diffusion studies (Maller et al., 2019). It was later reinforced that the hippocampus had several direct connections with cortical regions, including strong connections with early visual cortical areas (Huang et al., 2021). In other words, the hippocampus might be communicating with cortical regions without a gateway region such as ERC. Indeed, anterior-posterior extent of the human hippocampus is preferentially linked to cortical regions (Dalton et al., 2022). The current processing pipeline developed for the hippocampal circuit can easily be adapted to investigate the connections of the hippocampal subfields with various regions.

Human hippocampal pathways have not been extensively studied *in vivo* despite the advancements in neuroimaging methods. One concern in investigating these anatomical connections using diffusion imaging is the abundance of crossing fibers in the human brain. Novel processing algorithms such as Constrained Spherical Deconvolution have been proposed to overcome these limitations. This algorithm estimates a distribution of fiber orientations in each voxel in a diffusion image, unpacking the information in the voxels regarding the crossing fibers and thus yielding a much more accurate tractography (Tournier et al., 2007). Incorporating such algorithms with high resolution images are proven to be effective in *in vivo* verification of human white matter connections of small and complex structures such as the hippocampus (Elsaid et al., 2022; Huang et al., 2021; Karat et al., 2023; Maller et al., 2019; Treit et al., 2018).

We leveraged a similar methodology to quantify and characterize white matter connections within the hippocampus, pioneering *in vivo* human investigations of hippocampal pathways. Although the diffusion data we are utilizing is multi-shell, the resolution of anatomical images might be a limitation. For example, we were not able to detect the white matter fibers between DG and ERC. Similar issue was encountered in a previous *ex vivo* diffusion tensor imaging study that visualized perforant pathway in the human hippocampus (Augustinack et al., 2010). High resolution diffusion and anatomical data with faster repetition time and multi-shell acquisitions could be implemented to tackle these limitations.

There were also certain limitations leading to relatively less connections in the core of the hippocampus although some posterior connections entered relatively deeper layers of the hippocampus. The quality of the segmentation implemented in the pipeline has a prominent effect on the resulting hippocampal pathways. It is highly recommended that high resolution T2 weighted images, especially in coronal slices, are acquired for higher specificity. As this was unavailable in the current dataset, the segmentation was performed on T1 weighted images using the Multiple Automatically Generated Templates for different Brains (MAGeT Brain) algorithm (Chakravarty et al., 2013). As part of the automated segmentation procedure within MAGeT Brain, we implemented the atlas of the Extra-Hippocampal White Matter Atlas and Hippocampal Subfields (Amaral et al., 2018; Winterburn et al., 2013). Within this atlas, the deeper layers of the hippocampus such as stratum radiatum/stratum lacunosum/stratum moleculare were labelled separately from the hippocampal subfields. Although such distinction has an anatomical significance, these layers span CA1, CA2 and CA3. In the current work, it was necessary to exclude these deeper layers to avoid any overlap between the segmented regions that were used to mask the whole brain tractographies. We provide a processing pipeline for quantification of human hippocampal circuit with specific recommendations. Certain factors should be taken into consideration while characterizing hippocampal microstructure (Karat et al., 2024).

Overall, by harnessing cutting-edge neuro-analytic techniques, we extended our understanding of the hippocampal pathways that is established in theoretical, *ex vivo* and animal work to *in vivo* human investigations. We characterized MSP-, TSP-related and output connections at the participant level and evaluated the individual differences in the human hippocampal circuit.

Participant specific white matter connections provided here offer a healthy young adult normative data, which can be a benchmark for future studies in relating the hippocampal pathways to cognition. This also opens the door for multimodal investigations within the Human Connectome Project with its extensive data that is well beyond what is reported here. In addition, the population based hippocampal circuit provides a template, that future studies can use it as a reference to reconstruct the human hippocampal pathways. Lastly, we provide a well-documented processing pipeline that can be modified and utilized in reconstruction and quantification of the human hippocampal pathways in various populations from clinical or aging to developmental studies. Thus, this project paves the way for understanding the hippocampal circuit in humans and its implications in cognition, aging and diseases.

## 4. Methods

### 4.1. Human Connectome Project Participants

We analyzed diffusion and anatomical MRI volumes of 831 healthy adult participants (F = 445, M = 386; ages 22 - 35) from the Human Connectome Dataset, specifically Human Connectome Project 1200 subject release (Van Essen et al., 2013). Out of 1206 subjects in this release, we included those who had at least one T1, one T2, and one diffusion weighted volume. Within this subset, 57 participants were reported with structural or segmentation abnormalities by the Human Connectome Project team, 65 resulted with a faulty segmentation, and 3 yielded problematic Fiber Orientation Distribution (FOD) images in the MRtrix pipeline, leaving a total of 831 subjects for the central analysis. All Human Connectome Project participants provided informed consent. The study was approved by the Institutional Review Board at Washington University in St. Louis. The use of the Human Connectome Project data for the current analyses was conducted in accordance with the University of Toronto Research Ethics Board.

### 4.2. Human Connectome Project MRI Acquisition

Details related to image acquisition (Sotiropoulos et al., 2016) and preprocessing (Glasser et al., 2013) are published elsewhere. Scans were performed in a 3T Siemens Connectome Skyra equipped with a 100 mT/m gradient set and 32-channel receive coils (Uğurbil et al., 2013).

Nominal spatial resolution was 1.25 mm isotropic (matrix size PE × Readout = 144 × 168 with left–right (LR) phase encoding (PE) and 6/8 PE partial Fourier). A total of 108 echoes are collected, with echo spacing of 0.78 ms and readout bandwidth 1490 Hz/pixel, resulting in a total echo train length (ETL) of 84.24 ms. Sampling in q-space includes 3 shells at b = 1000, 2000 and 3000 s/mm2 and 18 b = 0 s/mm2 images. TE (89 ms) and TR (5.5 s) are matched across shells.

Total scanning time for this protocol was about 55 min. The Human Connectome Project minimal preprocessing pipeline used on this dataset (Glasser et al., 2013) included artifact removal, motion correction, and registration to standard space.

### 4.3. Hippocampal Subfield Segmentation

The Multiple Automatically Generated Templates for different Brains (MAGeT Brain) algorithm (Chakravarty et al., 2013) was used to segment the hippocampus into its subfields. This segmentation was performed using the Extra-Hippocampal White Matter Atlas and Hippocampal Subfields (Amaral et al., 2018; Winterburn et al., 2013) that generated the following subfields separately in left and right hemisphere; CA1, subiculum, CA4/dentate gyrus, CA2/CA3, as well as stratum radiatum/stratum lacunosum/stratum moleculare which was not included in the tractography.

This procedure also produced hippocampal white matter segmentation, including fimbria, mammillary body, fornix, and alveus. White matter segmentations were used as a mask to increase the specificity of TSP-related connections. MaGet Brain automates a multi-atlas-based segmentation procedure in which a subset of participants is first registered to manually labeled atlases to generate a library of template segmentations. Each participant from the entire dataset is then registered to each of the template participants and the multiple related template segmentations are then combined via voxel-by-voxel label fusion to generate a final participant-specific segmentation. For datasets with only T1 anatomical volumes, the multi-atlas segmentation approach has been demonstrated to be more sensitive and accurate relative to other single-atlas methods (Bussy et al., 2021; Chakravarty et al., 2013; Yushkevich et al., 2014).

### 4.4. Entorhinal Cortex Segmentation

We segmented ERC, using the Automatic Segmentation of Hippocampal Subfields (ASHS) algorithm (Yushkevich et al., 2014) and ASHS Penn Memory Center T1-Only Atlas (PMC-T1) (Xie et al., 2016). This segmentation process also included the medial temporal lobe (MTL) regions (i.e., perihinal and parahippocampal cortices along with ERC) and anterior/posterior hippocampus. ASHS first creates an atlas library of manually segmented brains each registered to a template participant. Then, labels are fused across atlases to generate the final segmentation. This procedure was completed using the Distribution Segmentation Service of ITK-SNAP (Yushkevich et al., 2006). Within the resulting segmentation, only ERC was combined with the hippocampal subfield segmentation and used in the remaining analyses.

### 4.5. Fiber Orientation Distributions

Whole brain tractography and resulting connectomes for each participant were generated (Fig. 6a), using MRtrix3 (J. D. Tournier et al., 2019). This tool allows for the characterization of fiber populations within each voxel (i.e., fixel) through Multi-Shell, Multi-Tissue Constrained Spherical Deconvolution (MSMT-CSD) and produces a precise estimation of crossing fiber orientation (Jeurissen et al., 2014; Tournier et al., 2012). We first estimated response functions for three different tissue types (white matter/gray matter/cerebrospinal fluid) based on co-registered five-tissue type (5TT) images derived from participants’ T1 images (*dwi2response* with *msmt_5tt* algorithm) (Jeurissen et al., 2014) (Fig. 6a). We then estimated fiber orientation distribution (FOD) based on these response functions (*dwi2fod* with *msmt_csd* algorithm) (Jeurissen et al., 2014).

**Fig. 6.**
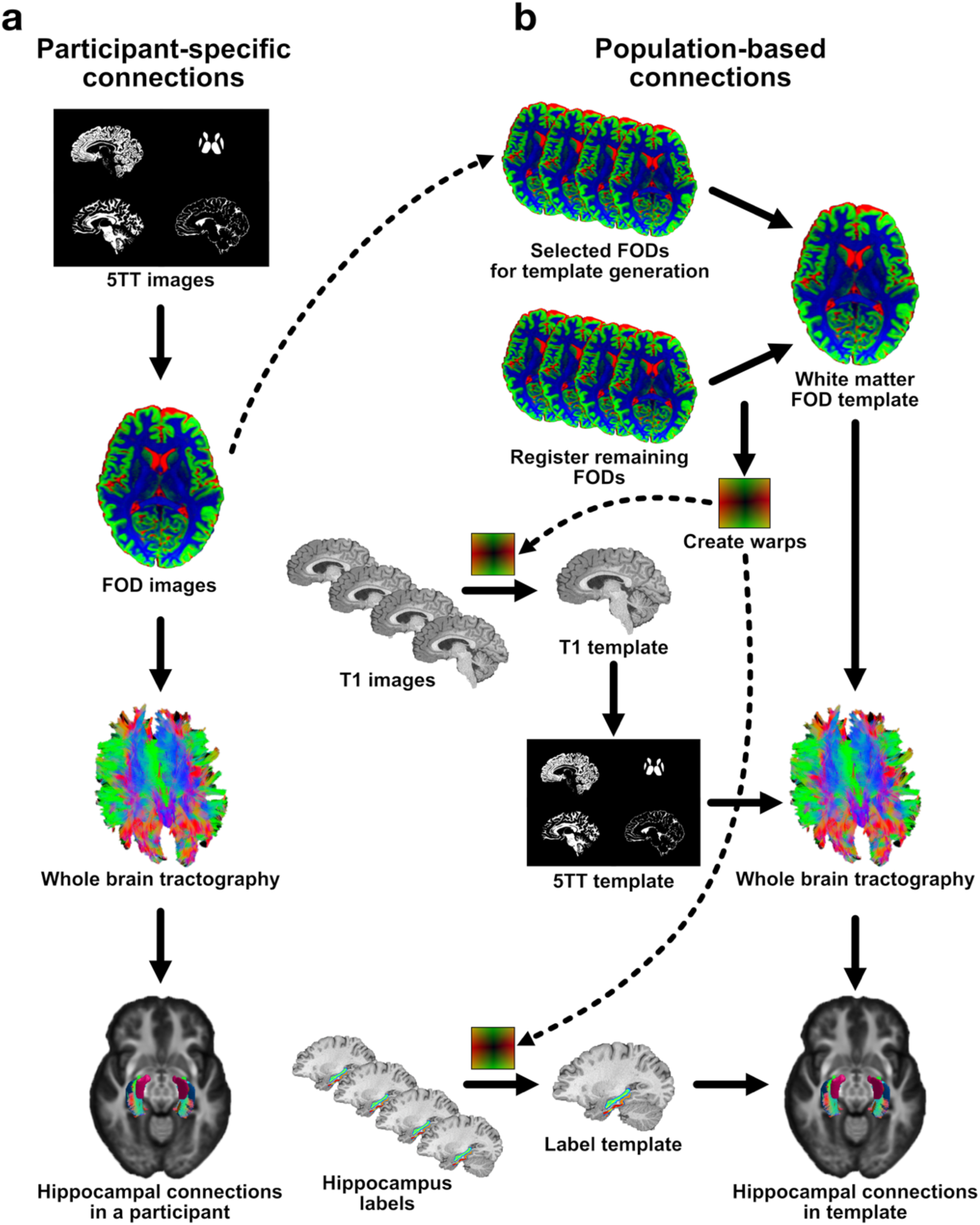
Processing pipeline for quantifying hippocampal white matter connections in humans. a, First arm of the processing pipeline is for reconstruction of hippocampal white matter connections at the individual level. The processing includes generation of 5 tissue type images, estimation of fiber distribution images, reconstruction of whole brain tractography and creation of hippocampal connections by masking the whole brain tractography by the segmentation images that include hippocampal subfields and ERC. **b,** Second arm of the pipeline enables the creation of hippocampal white matter connections based on the entire sample. This follows the same steps as the first arm; however, the processing here is based on the template images that are warped into the template space, not subject-specific images.

### 4.6. Whole-Brain Tractography Reconstruction

Based on the white matter FOD images, whole-brain probabilistic tractography for each participant was created using *tckgen* with the *iFOD2* algorithm and the Anatomically-Constrained Tractography (ACT) framework (Smith et al., 2012; Tournier et al., 2019). Each tractography was generated with 10 million streamlines, using the curvature threshold of 45°, maximum fiber length of 250 mm while seeding from the gray matter-white matter interface and retracing poor structural terminations (Smith et al., 2012; Tournier et al., 2010). We then applied spherical deconvolution informed filtering to match streamline densities with the FOD lobe integrals, allowing the removal of false-positive tracks and creating more biologically meaningful estimates (Smith et al., 2013). With this step, final whole-brain tractography of each participant included 2 million streamlines.

### 4.7. Human Hippocampal Pathways at the Individual Level

Structural connectomes were generated based on subject-specific whole-brain tractography and segmentation image (hippocampal subfields + ERC) using *tck2connectome* (Tournier et al., 2019). Each resulting connectome contained the density information of streamlines between the hippocampal subfields and ERC. Each individual hippocampal connectivity matrix consisted of 5 regions of interests (ROIs) in each hemisphere: CA1, subiculum, CA4/dentate gyrus, CA2/CA3, and ERC. Each entry in these connectivity matrices represents the streamline density between 2 ROIs in each participant specific segmentation image.

Based on the resulting connectomes, we quantified the streamlines for the theorized hippocampal pathways: 1) TSP-relate connections refer to DG (combined with CA4) - CA3 (combined with CA2) and CA1-CA3, 2) MSP-related connections include ERC-CA1, 3) output connections include those between CA1-SUB and SUB-ERC. Unfortunately, we were unable to track white matter connections of ERC-DG, which is part of TSP. The difficulty in detecting these fibers have been previously reported (Augustinack et al., 2010). DG has a deep convolution and significantly small size (Hevner, 2016), which might make it difficult to identify the fibers between ERC and SUB. This might be due to the curvature of the hippocampus such that SUB is located between ERC and DG, and thus, most of the connections between ERC and DG might be instead terminating at SUB in our analyses (Augustinack et al., 2010).

Characterizing participant specific hippocampal pathways allowed us to estimate the proportion of connections among the hippocampal subfields and ERC, indicating the relative streamline density of such connections. The streamline density of these connections was further explored in terms of hemispheric and sex differences. We leveraged the individual variability within the hippocampal connections to explore their relation to various cognitive measures.

### 4.8. Population Template of White Matter Fiber Orientation Distributions

White matter FOD images of 30 participants, representing the available sex and age range, were selected for population template creation (Fig. 6b). Remaining participants’ FOD images were then registered to this white matter FOD template. Warps created during this registration were used on both individual T1 and segmentations (hippocampal subfields + ERC) to align and orient them in the population template space. Warped T1 images were averaged for creating the T1 template, which was then used to create the 5TT template. A label template was generated by aggregating across participants’ warped label volumes and selecting for each voxel the ROI label observed in the majority of participants. Lastly, the 5TT and white matter FOD templates were used to create whole brain tractography for the population. The label template was applied to the tractography to create a population-based connectome which was used to identify white matter streamlines among the hippocampal subfields and ERC. Streamlines between each ROI pair were edited using exclusion masks which consisted of 2.5 – 5 mm dilated ROI labels.

### 4.9. Quality Control

All T1 images included in this study were visually inspected by M.G. and A.B. Participants flagged with abnormalities overlapped with other exclusion criteria and were excluded. In addition, we computed the Pearson correlation between the individual T1 images and the T1 template. Approximately 96% of participants yielded a correlation coefficient equal to or larger than 0.85, showing a reliable match between individual T1 images and the template created.

Similarly, subject specific hippocampal subfield and ERC segmentations showed a reliable overlap with the label template as 99% of participant segmentations yielded a DICE coefficient of ≥ 0.5. ROI volumes were also calculated and corrected with intracranial volume. Streamline densities of hippocampal pathways were corrected for the ROI volumes. However, uncorrected streamline densities are reported here because volume correction did not make a significant difference in streamline distributions.

### 4.10. Relating Human Hippocampal Pathways to Cognition

The hippocampus is involved in variety of cognitive processes including learning, memory, and spatial navigation. We investigated the relationship between the human hippocampal pathways we quantified to the cognitive assessments available within the Human Connectome Project. We selected 5 tasks that tap onto episodic memory, working memory, and spatial orientation, as well as card sorting (Corcoran & Upton, 1993) that were shown to involve the hippocampus. Thus, we included picture sequence memory score (i.e., episodic memory), Penn word memory correct responses (i.e., verbal episodic memory), and list sorting score (i.e., working memory), variable short Penn line orientation total correct (i.e., spatial orientation), and dimensional change card sorting score (i.e., cognitive flexibility). For each cognitive assessment, we built separate linear mixed effects models to estimate streamline density of participant-specific hippocampal pathways with a fixed effect of cognitive score and random effect of individual differences. Age and biological sex were included as covariates. We corrected for multiple comparisons using the Bonferroni Test with an alpha level of 0.01 (α = 0.05/5 cognitive tasks).

## 5. Code and Data Availability

Our complete processing pipeline including specific codes and resulting data will be available at https://github.com/macklab/hippcircuit upon publication. For requests prior to publication, please

contact the corresponding author, Melisa Gumus, at.

## Acknowledgements

This project was funded by Natural Sciences and Engineering Research Council (NSERC) Discovery Grants (RGPIN-2017-06753 and RGPIN-2024-0588 to MLM), Canadian Institute of Health Research (CIHR) Grant (PJT-178337 to MLM), Brain Canada Foundation Grant (to MLM), and Vanier Canada Graduate Scholarship provided by NSERC (to MG). This project was only possible through the additional computing allocations awarded by SciNet high performance computing consortium (Compute Canada). We would like to thank the CobraLab at the McGill University for giving us access to MAGetT Brain on Compute Canada. We would also like to thank Bradley Karat at the Western University, Robert Smith at the University of Melbourne, and Ingrid R. Olson at the Temple University for insightful analyses suggestions.

## 6. Author Contributions

M.G. conducted the analyses, developed the processing pipeline, and wrote the manuscript. A.B. and M.G. developed and built the website associated with the project. M.L.M. supervised the project and directed project progress. All authors edited the manuscript.

