## Extended Data Figure 1 for "*In vivo* Quantification of White Matter Pathways in the Human Hippocampus"

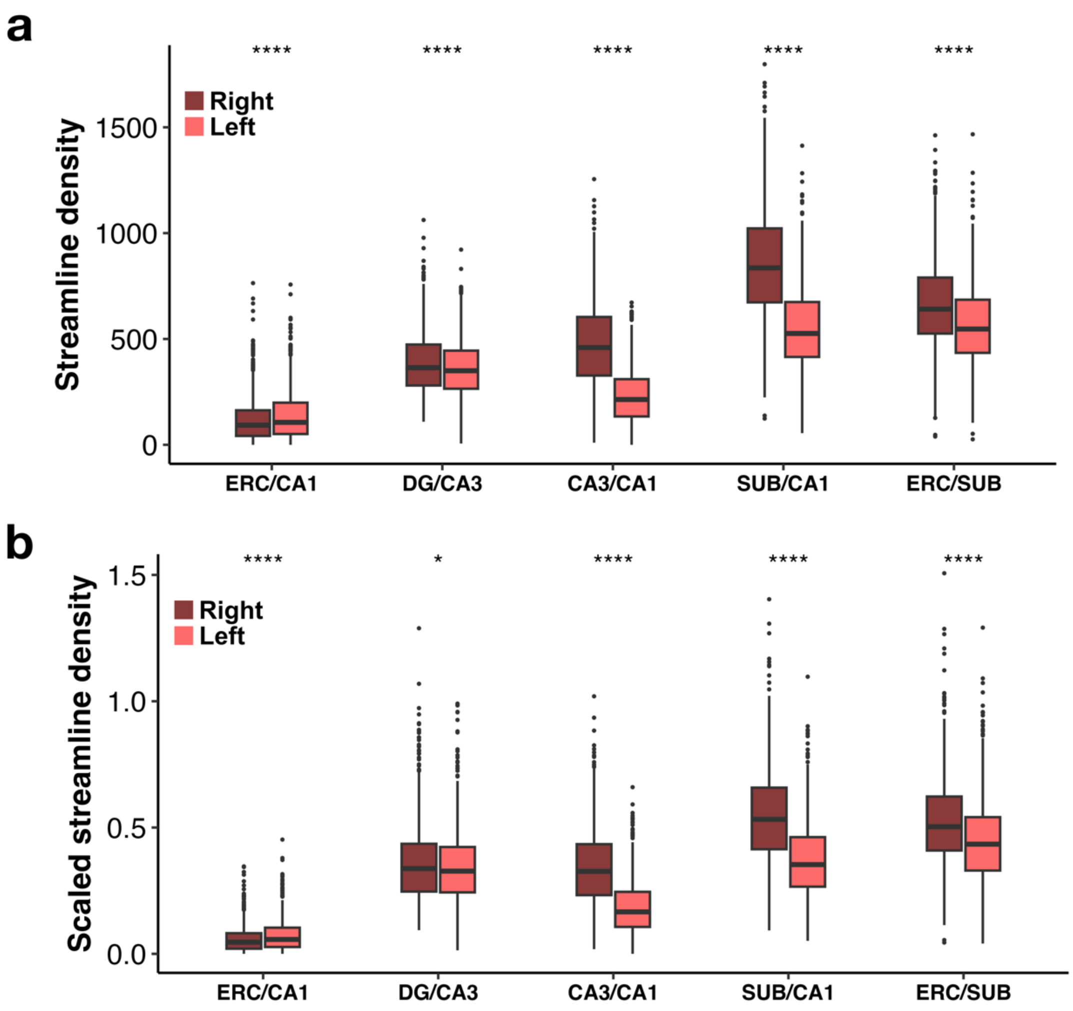


**Extended Data Fig. 1: Hemispheric differences in streamline density of the quantified human hippocampal pathways. a,** MSP-related connections present greater streamline density in the left hippocampus while TSP-related and output connections are more prominent in the right hemisphere. **b,** Streamline density of each hippocampal pathway is corrected by the average volume of the 2 regions it connects to. Volume differences across the subfields and ERC did not influence relative streamline densities.
